# Histopathological Study Of Different Organs Of Charles Foster Rats Under The Exposure Of *Pueraria tuberosa*

**DOI:** 10.1101/671529

**Authors:** Harsh Pandey, Shivani Srivastava, Mohan Kumar, Yamini Bhusan Tripathi

## Abstract

The present study was undertaken to investigate the safe doses of Pueraria tuberosa water extract (PTWE) on different organs. The OECD guidelines 407 of repeated toxicity was followed with respect to the selection of dose and days for different organs. The selected doses of PTWE were 250, 500, 1000 and 2000 mg/kg b wt for 7, 14, 21 and 28 days. Haematoxylin and eosin staining was used to study the morphological alterations in heart, intestine, testis, adrenal gland and spleen. In the present study, no adverse alterations in cardiac fibers of the heart, size and shapes in crypts and villi of intestine, seminiferous tubules and spermatozoa count in testis, three zones of adrenal gland, and spleen were seen in all treated groups of PTWE. There were no adverse morphological alterations found in described organs. The PTWE are safe at 1000 mg/kg b wt. up to 28 days and 2000 mg/ kg b. wt up to 21 days, respectively.

## INTRODUCTION

In several countries, more than 60% of the population are dependent on herbal medicine for health care (Saminathan *et al.* 2013). However, because of non-availability of safety parameters, many people are hesitant in using herbal medicine. Although it is well established that herbal medicines have therapeutic responses and they are already in clinical use (Saminathan *et al.* 2013).

The powder of *Peuraria tuberosa* (PT) tubers is in clinical use in Ayurveda as health promotion medicine (Sherman *et al.* 2010). In Hindi, it is called as vidarikand and in English it is known as Indian kudzu. It belongs to the family Fabeacea (Prasain *et al.* 2012). Its major secondary metabolites include puerarin, tuberostan, genestein, daidzin, tuberosin, puerarin, and pterocarpan puerarone (Rastogi *et al.* 2013, Maji *et al.* 2014a,b). In kudzu root, puerarin is the most abundant (approx. 23%w/w) and it has potents ability to cause various pharmacological effects (Lee *et al.* 2005). *Pueraria tuberosa* has shown many pharmacological activities in rats (Pandey *et al.* 2020). Pharmacokinetic study of PT was studied by Pandey *et al.* (2019b).

Recent past, it has been that *Peuraria tuberosa* water extract (PTWE) act as antidiabetic herbal drug working through incretin signalling pathway (Srivastava *et al.* 2018). Further regarding its mechanism of action towards anti-diabetic and nephroprotective potential by inhibition of DPP4, MMP-9, PKC-beta, and activation of Caspase enzymes. The results have shown activation of SOD, catalase, BCL-2, and nephrin (Srivastava *et al.* 2015, Srivastava *et al.* 2017, Shukla *et al.* 2016). However, for the first time we reported its action of GLP-1, GIP, TNF-alpha, IL-6 and HIF-1 which were inhibited by *Pueraria Tuberosa* in different experimental conditions such as diabetes and kidney damage (Srivastava *et al* 2019). These results are according to earlier reports related to the role of different pure phytochemicals, which are also found in PT tubers (Bebrevska *et al.* 2007), (Cai *et al.* 2011).

The histopathological examination is the golden standard for evaluating treatment related pathological changes in tissues and organs (OECD 1995). The pathological study can add information to the clinical data. Histopathological study is very clear, reliable parameter to study the effectiveness/toxicity of drugs in animal tissues. Study of safety and toxicity are the main parameters used for before the clinical use of drugs.

Earlier, were ported the effect of PTWE in pre-clinical toxicity in rats as per OECD guideline 425 and 407 with special reference to changes in liver and kidney functions (Brondani *et al.* 2017, Pandey *et al.* 2018a). In the present study, partially purified water extract of tubers of *Peuraria tuberosa* (PTWE) has been used to assess its toxic effect on Charles Foster rats as per OECD guidelines. The herbal tablets of PTWE were prepared by a wet granulation method for clinical study (Pandey *et al.* 2018b). In the present study, effect of *Peuraria tuberosa* on other organs after oral treatment with different doses for different durations up to 28 days. The histological changes have been observed in heart, intestine, testis, adrenal gland and spleen in rats of Charles foster strain.

## MATERIALS AND METHODS

### Collection of plant material and preparation of extract

The plant material *Pueraria tuberosa* (PT) was purchased from Ayurvedic Pharmacy BHU Varanasi, preserved in our laboratory, Department of Medicinal Chemistry IMS BHU (Ref. no YBT/MC/12/1-2007) and also with the sample preserved in Museum of Department of Dravya-guna, of our Institute IMS BHU. It was further compared with Thin Layer Chromatography finger printing with standard sample, which has been characterized earlier in our laboratory Department of Medicinal Chemistry by DNA fingerprinting, High Performance Thin Layer Chromatography and Light Chromatography-Mass Spectrometry (Nagwani *et al.* 2010). Pharmacognostic characterization of crude powder of PT done on the basis of standard pharmacopeia (Pandey H *et al.* 2019a). The water extract of plant *Pueraria tuberosa* was prepared by water decoction method. The yield value of extract was 36% w/w. It was characterized by TLC finger printing.

### Animals

Charles Foster strain of rats were purchased from the Central Animal House of our Institute IMS BHU (542/GO/ReBi/S/02/CPCSEA). The animals were acclimatized for 7 days in laboratory condition and subjected to anti-protozoa treatment by giving drug Metronidazole orally. Finally, the animals were randomly divided into five groups of 6 animals in each. The experimental protocol was approved by the Institutional Ethical Committee (Dean/2017/CAEC/721).

### Experimental design

The selection of dose and time of drug treatment was followed as per earlier studies and the experiment was carried out as per OECD guidelines 407 (Brondani *et al.* 2017).The dose was prepared by dissolving PTWE in water with gum acacia and different dose (250 mg/kg b. wt, 500 mg/kg b. wt, 1000 mg/kg b. wt, and 2000 mg/kg b. wt) were orally given for 28 days, to each rat of the respective groups, in the morning time. Weekly assessments of body weight and diet intake were recorded. At the end, the animals were sacrificed by anesthetizing the rats by intra-peritoneal injection of pentobarbitone sodium (45 mg/kg b. wt). The blood was collected within Immediately of death, all the required organs were dissected out. The attached fat and other undesired attached tissues were dissected out and finally the organs were drained on the blotting paper and weighed on electronic pan balance. The chosen organs were heart, intestine, testis, adrenal gland and spleen, which were finally fixed in formalin for histopathological study. The animals of different groups were sacrificed at different time intervals of 7, 14, 21 and 28 days. The tissues of the group having a dose of 2000 mg/kg b. wt could not be collected at 28 day, because all animals of this group died.

### Histopathology

Organs such as heart, spleen, testis, intestine, and adrenal gland were isolated and trimmed of excess fat in each group of animals, fixed in formalin for H & E staining. The standard protocol of making micro sections and staining was adopted (Troyer 2008). The fixed tissues were taken out of the fixative, washed properly and small sections were processed for dehydration and finally embedded in the paraffin wax. After solidification, the blocks were trimmed, mounted on microtome (medimeas/mrm-1120 A) and thin sections of 5-6 μm thickness were cut and placed on slides coated with albumin and then finally subjected to a process of dewaxing, dehydration and staining with haematoxylin and eosin (H &E) staining. Finally the sections were mounted under a cover slip and sealed with Dopex Mount. The Transverse Sections of all the organs were examined and image were taken with binocular fluorescent microscope without lamp on (Nikon Eclipse 50i Japan), fitted with a digital camera. Histological analysis was done to further confirm the alteration in cell structure of the organs. All the slides were randomly photographed in 10 view fields. The measurements of captured photographs were done by using software NIS Elements Basic Research.

## RESULTS

The Gross lesions in heart, intestine, testis, adrenal gland and spleen were found to be normal as earlier reported. Most of them showed a high degree of liver toxicity, as already reported in our preclinical toxicity study (Pandey *et al.* 2018a).

### Control animals

The heart tissue section has normal intact cardiac muscle myofibrils and intercalated disc (Fig. 1A). In the intestinal tissue, glands, mucosa, sub-mucosa, serosa, muscularis externa and villi are normal (Fig. 1B). The spermatogonia, spermatozoa, and spermatids are normal in testis (Fig. 1C). Adrenal gland showed normal an outermost portion cortex and inner section medulla (Fig 1D). Splenic section showed normal red and white pulp areas (Fig. 1E).

**Fig. 1.**
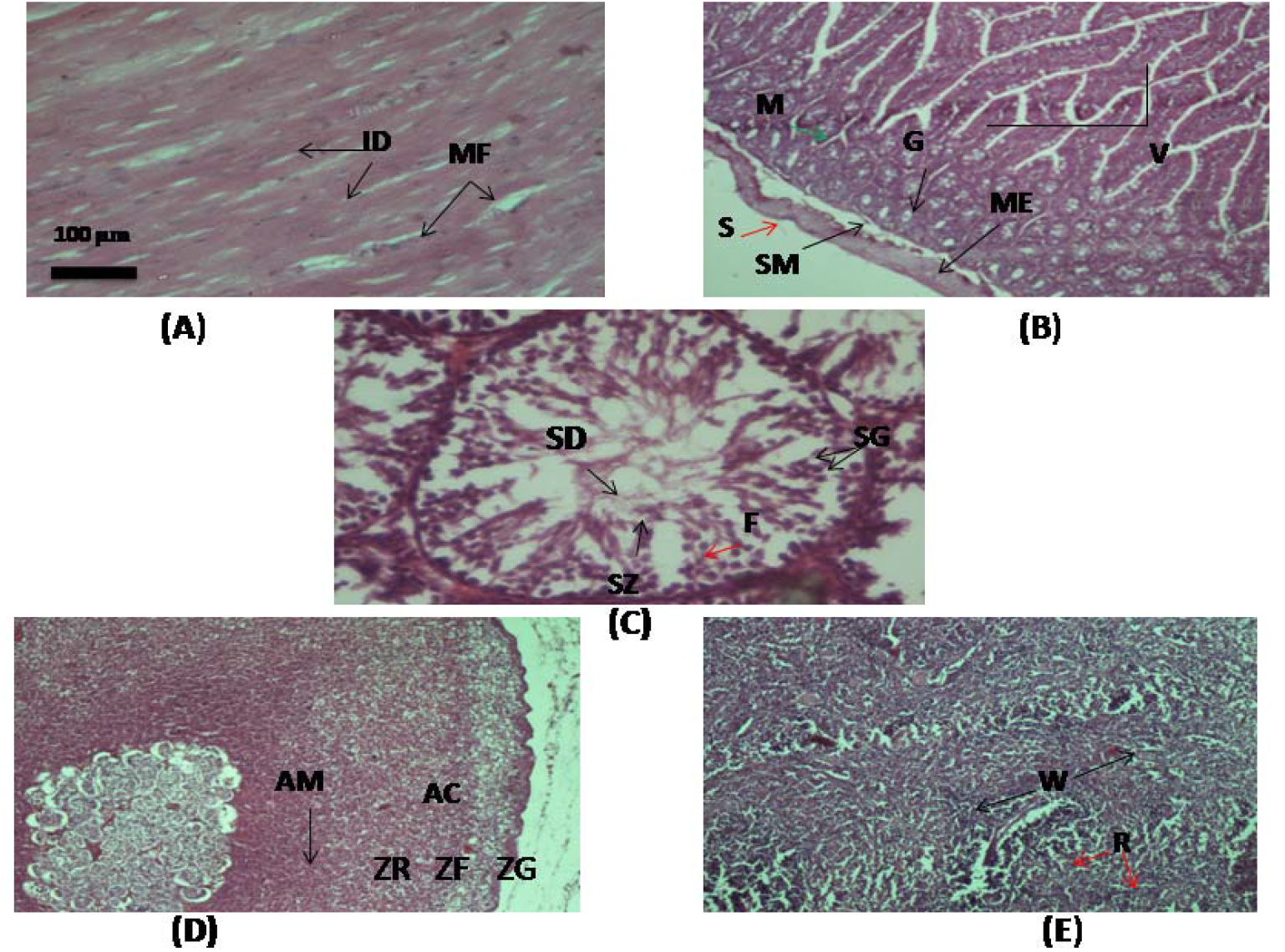
Histology of normal rats (A) Heart, (B) Intestine, (C) Testis, (D) Adrenal gland and (E) Spleen. [Scale bar 100 μm with different magnification like 10x and 40X according to specific organs site. ID-Intercalated disc, MF-Myofibrils, G-Glands, M-Mucosa, SM-Sub mucosa, S -Serosa, ME-Muscularia externa, V-Villi, SG-Spermatogonia, SZ-Spermatozoa, SD-Spermatids, F-Follicles, AM-Adrenal medulla, AC- Adrenal cortex, ZR-Zona glomerulosa, ZF-Zona fasciculata, ZG- Zona reticularis, R-Red pulp, W-white pulp].

### Effects of PTWE on different organs at different doses

#### Heart

The T.S. of heart did not show any changes in the extract treated group for all tested doses up to 21 days. The sections showed intact cardiac muscle myofibrils and intercalated disc (Fig. 1A). The treatment for 21 days with 2000mg/kg b wt also did not show any cellular damage or focal necrosis of cardiac muscle fibers. No interstitial oedema or swelling was also found (Fig. 2)

**Fig. 2.**
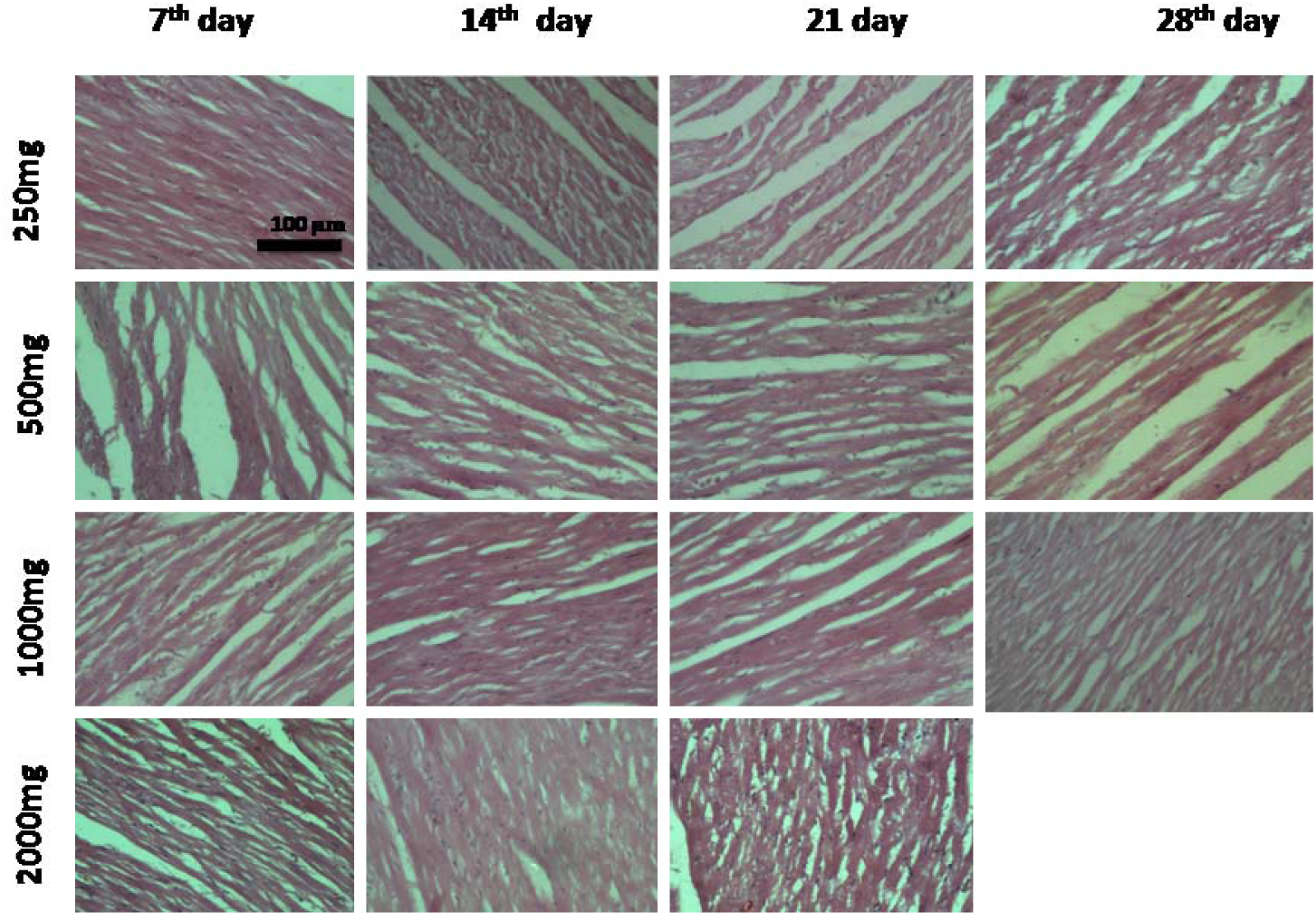
Histology of PTWE treated heart tissue at different time intervals with variable dosage. Scale bar 100 μm.

#### Intestine (Jejunum)

The intestinal part, which was selected for study was 10 cm distal to the duodenum. Jejunum showed no significant changes in the PTWE treated rats. The T.S. images were similar to normal rat intestine (jejunum) T.S. The serosa and mucous membranes were normal. The number and size of Villi were also normal. The glands found in the lower portion of intestinal wall were also normal (Fig. 3).

**Fig. 3.**
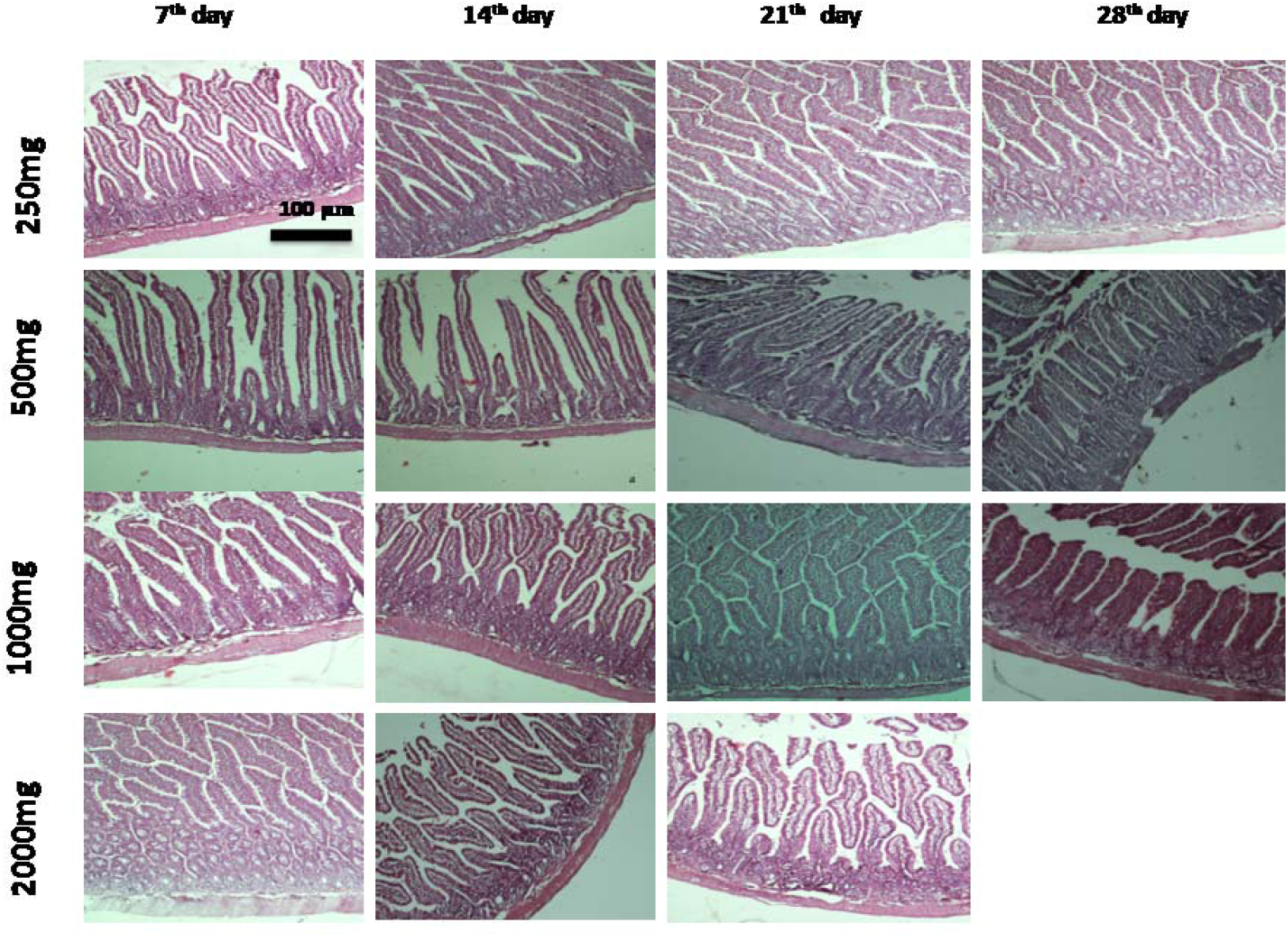
Histology of PTWE treated intestinal tissue at different time intervals with variable dosage. Scale bar 100 μm.

#### Testis

Histopathological examination of the testis showed normal histological structure of active mature, functioning seminiferous tubules associated with complete spermatogenic series are shown in (Fig.1 and Fig.4). The spermatozoa and spermatogonia was found in fully mature condition in all sections of tissue of doses up to 2000mg/kg b wt for 21 days. There were no congestion, edema and dilation were found in blood vessels (Fig.4). It may be effective in patients, already having a defective physiology.

**Fig. 4.**
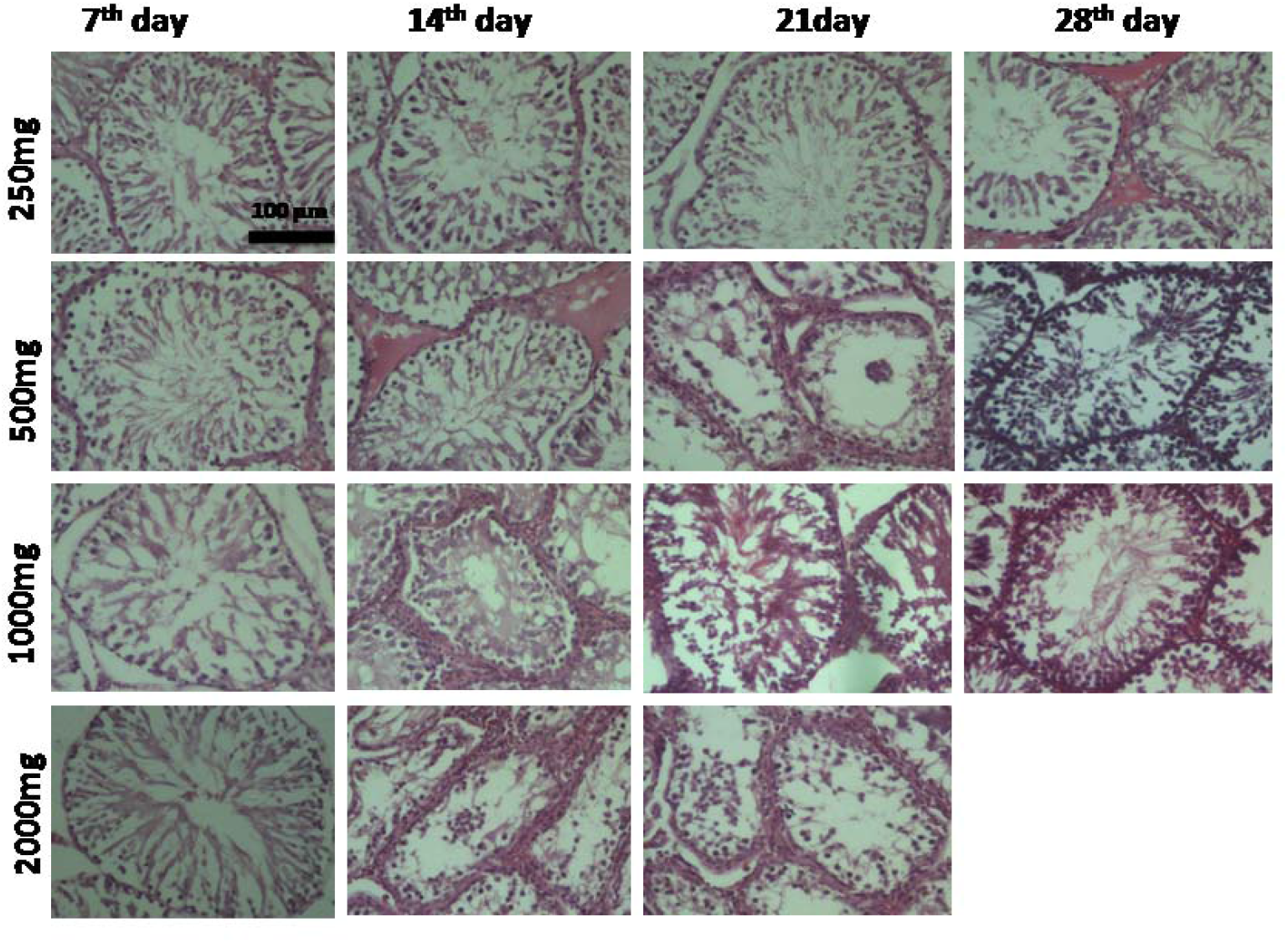
Histology of PTWE treated testis tissue at different time intervals with variable dosage. Scale bar 100 μm.

#### Adrenal gland

The zona glomerulosa, zona fasciculata, zona reticularis and adrenal medulla were intact in all tissue sections. No morphological dissimilarity found in the cortex region of the adrenal gland There is no amorphous debris was found in focal areas of zona fasciculata of the higher dose of PTWE *i.e*; for 100 and 2000mg/kg bw up to 21 and 28 days respectively (Fig.5).

**Fig. 5.**
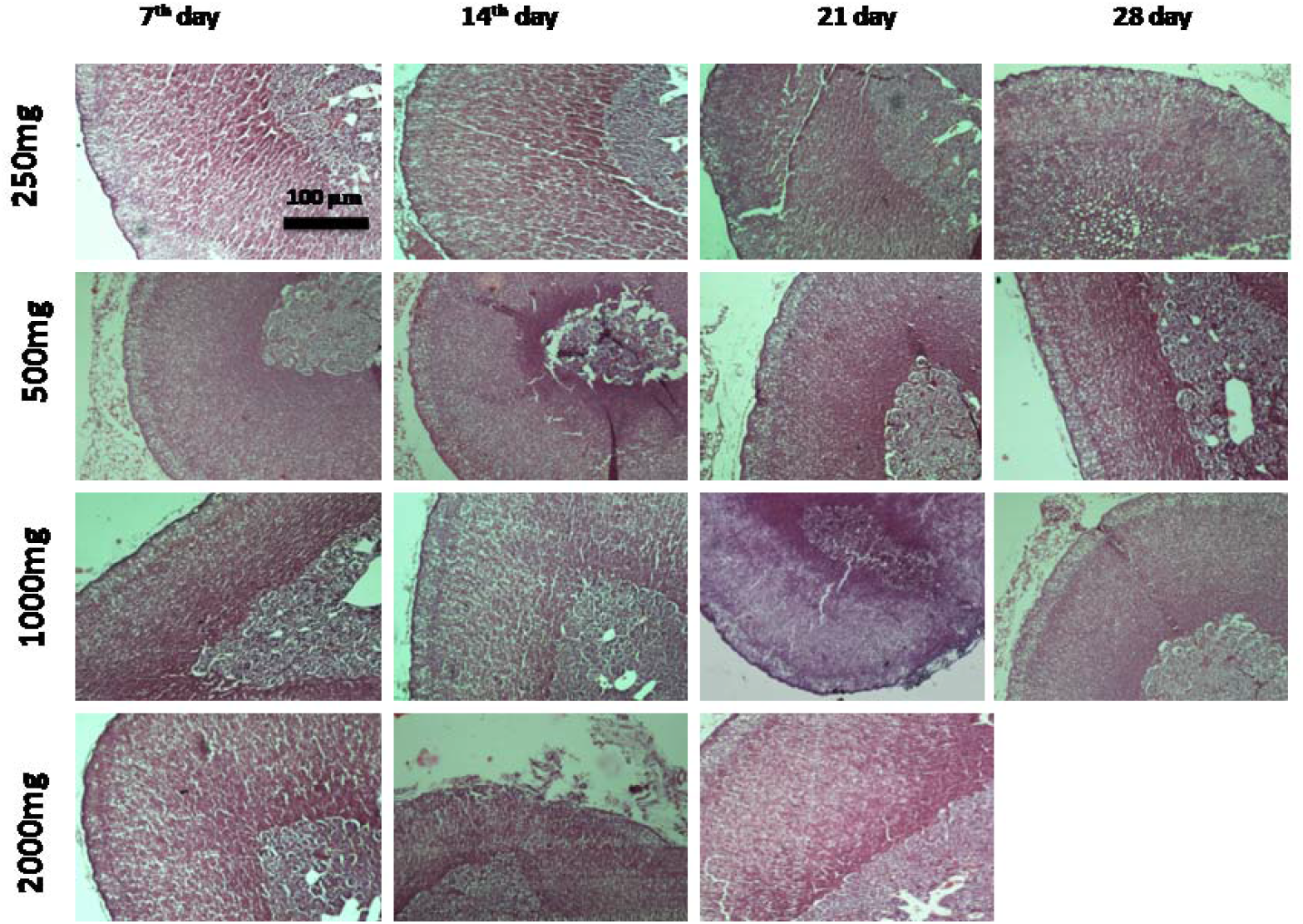
Histology of PTWE treated adrenal tissue at different time intervals with variable dosages. Scale bar 100 μm.

#### Spleen

The T.S. of spleen showed the normal architecture in red pulp and white pulp regions in all the tested doses of PTWE. The red pulp region consists of red blood cells, lymphocytes and plasma cells, whereas the white pulp region contains T lymphocytes and B lymphocytes. observed in the red pulp region was also normal (Fig.6).

**Fig. 6.**
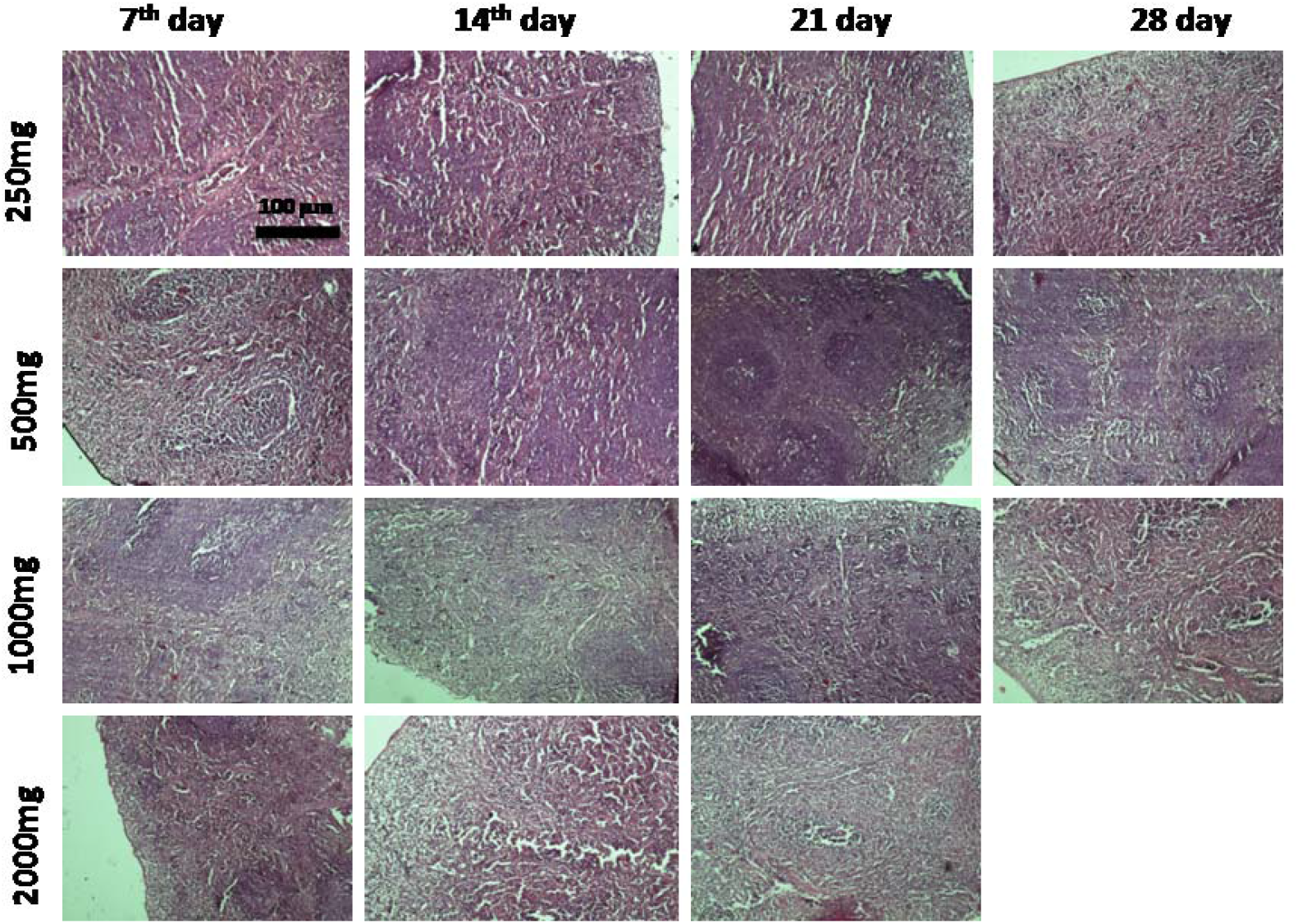
Histology of PTWE treated splenic tissue at different time intervals with variable dosages. Scale bar 100 μm.

As compared to growing the number of herbal drug users around the globe, there is a lack of scientific data on the safety profile of herbal products (Saad B *et al.* 2006), therefore the safety of these products have become an important issue (WHO 2004). In this experiment, we have conducted OECD guidelines 407 and found that histopathological examination of the vital organs did not reveal any morphological changes after oral administration of *Pueraria tuberosa* for 28 days at the dose level of 250, 500, 1000 and 2000 mg/kg bw.

## DISCUSSION

### Heart

In the present scenario, cardiovascular diseases, particularly, become a worldwide health problem affecting all economic groups of the society.

Tannins, flavonoids and glycosides have significant antioxidant properties, thus augments antioxidants and induction of HSP 72(Nieto *et al.* 1993). In the previously reported isoflavonoids and glycosides are found in plant *Puerraia tuberosa*. PT is also reportedly known for its antioxidant property (Pandey *et al.* 2007).The major bioactive component of the PT is flavonoids known as puerarin. On the basis of previous study of puerarin and our present result, we can assume peurarin might be the possible component of the PT which protect the cardiovascular tissue via a different mechanism as puerarin lower the mRNA & protein level and its receptor APJ in one clip hypertension (Jin *et al.* 2009), and blocks TSP 1 expression in diabetic rats (Pan *et al*. 2009). s

### Intestine

The Intestine is the main primary organ, responsible for food materials and drugs absorptions. The small intestine comprises 4 layers, mucosa, sub mucosa, muscularis externa and serosa. Mucosa consists of 3 layers, epithelium, lamina propria and muscularis mucosa organized into villi and crypts (crypts of lieberkuhn). Villi are finger like projection of the epithelium which contain blood and lymphatic vessels into the intestinal lumen. Intestinal mucositis is a common side effect of clinical chemotherapy for patients with cancer (Wadler *et al.* 1998), and includes symptoms such as severe diarrhoea and dehydration. The anti metabolite anticancer agent, 5-fluorouracil (5-FU), is widely used to treat several types of malignant tumours, it frequently causes intestinal mucositis. Mucosistic is morphologically characterized by the shortening of villus height and destruction of crypts in the small intestine. Previously the apoptosis is detected in intestinal crypts 24 hours after the first administration of 5-FU in mice. Those puerarin responsible for anti-inflammatory activities via inhibiting the level of IL –8(Pang *et al.*, 2012) and suppressed the protein or the mRNA expression of TNF – alpha, NF-kB, iNOS, TGF- B1 and MDA (Li *et al.* 2013). In our result, no significant adverse effect was found in different doses of *Pueraria tuberosa* on morphological dissimilarity in size and shapes of the villi and crypts of intestine since it was reported that PT have more isoflavonoids.

### Testis

The results were contrary to the literature, showing its anti-infertility and spermatogenic activating potential, at least in normal rats. It may be effective in patients, already having a defective physiology.

Drug induced toxicity was found in the testis by cisplatin and metabolic disorder such high fat diet induced diabetic model in rats. Various types of isoflavones, phyto-estrogens, puerarin etc. may be responsible for the stimulation of androgenic activity has been reported in the roots of *Pueraria tuberose* (Chauhan *et al.* 2011, Chauhan *et al* 2012). Phytoestrogen like daidzein and genistein also affect neurobehavioral are anti-oestrogenic either an action in opposition to that of oestradiol and increase the level of LH, FSH, and testosterone via stimulation of gonadotrophin releasing hormones GnRh (Lee *et al.* 2002, Patisaul 2005). Kudzu and puerarin combination with p4 may synergistically interfere with NADPH oxidation was reported in chromatin condensation (Chapman and Michael 2003, Bennetts *et al.* 2008). In our results no any types of morphological dissimilarities were found in rats after 28 days treatment of PTWE.

### Adrenal

We know that in adrenal gland necrosis found more frequently in the cortex (especially zona fasciculata and reticularis) than medulla. Flavonoids like Daidzein and nobiletin those found in PT was earlier reported for less, but significant increase in catecholamine synthesis or secretion via activation of extracellular signal regulated protein kinases (ERKs) through the plasma membrane estrogen receptor (Liu *et al.* 2007) thus enhance the symapatho-adrenal system. The histological changes in adrenal could be correlated to its function. It is already reported that the release of the corticosteroids cortisol and aldosterone can be stimulated through the sympatho-adrenal system, by mediation through chromaffin cells in a paracrine manner. But here no such changes have been observed in our study, suggesting no any types of adverse effect of PTWE on these systems.

### Spleen

The isoflavones like puerarin, daidzein, and genistein are the most important constituents, and these are responsible for the immunomodulatory function (Sawale *et al.* 2013). The effect of *Pueraria tuberosa* and its isoflavones on haematopoietic system as well as on the function of T cells and neutrophils have been extensively correlated with immunomodulatory function.

The leukocytes including neutrophils, lymphocytes, monocytes, eosinophils, and basophils are responsible for immune response. In our experimental study histo-pathological investigation of spleen did not exhibit any abnormalities treated with a low or high dose of Pueraria extract. Also, spleen appeared grossly coloured and no finding of congestion was found in all treated tissues. The tissue is clearly differentiated into white pulp (WP) and the red pulp (RP). The architecture of the white pulp displayed normal rounded scattered follicles.

## CONCLUSION

No adverse alterations were founded in cardiac fibers in the heart, size and shapes in crypts of intestine, seminiferous tubules and spermatozoa were normal in testis. Three zones of adrenal gland were normal, and no adverse changes were seen in the pulps of spleen in all the groups treated with 2000 mg /kg b. wt of PTWE up to 21 days and with 1000 mg/kg b. wt of PTWE up to 28 days. After completed that experiments found to be safe up to 2000mg/kg b. wt. treatment of PTWE after 21 day.

## AUTHOR’S CONTRIBUTION

HP and YBT planned the experiment. HP has done all the experiment and documentation. MK helped in Histology. HP, SS, and YBT wrote the manuscript.

## ACKNOWLEDGEMENT

We are highly thankful to Dr Shashikant Patne, Dr. Radha Chaubey and all lab technicians of Department of Pathology, IMS, BHU for their kind help in histology and also for providing facilities.

